# Preliminary genetic assessment of three goatfish species in the Mediterranean Sea

**DOI:** 10.1101/2021.05.23.445364

**Authors:** Taha Soliman, Joseph D. DiBattista, Reda M. Fahim, James D. Reimer

**Affiliations:** Okinawa Institute of Science and Technology Graduate University, Onna-son, Okinawa 904-0495, Japan; College of Fisheries and Aquaculture Technology - Arab Academy for Science, Technology and Maritime Transport (AASTMT), Alexandria 1029, Egypt; Molecular Invertebrate Systematics and Ecology Laboratory, Department of Biology, Chemistry and Marine Sciences, Faculty of Science, University of the Ryukyus, Nishihara, Okinawa 903-0213, Japan; Australian Museum Research Institute, Australian Museum, 1 William St, Sydney, NSW 2010, Australia; School of Molecular and Life Sciences, Curtin University, Perth, WA 6102, Australia; Tropical Biosphere Research Center, University of the Ryukyus, Nishihara, Okinawa 903-0213, Japan

**Keywords:** Aswan High Dam, dispersal, Egypt, life history, mtDNA, Mullidae, salinity, sea surface temperature

## Abstract

We examined genetic diversity and connectivity of two indigenous Mediterranean goatfish species (*Mullus barbatus* and *M. surmuletus*), and a Lessepsian migrant species (*Upeneus moluccensis*), across the Nile Delta outflow using two mitochondrial DNA markers (COI and cyt b). Genetic diversity was high for the two Mediterranean species but relatively lower for the migrant species, suggesting founder effects after invasion from the Red Sea. Confirmation of this hypothesis, however, would require comparison with populations of origin in the Red Sea and the Indo-West Pacific. AMOVA and network analyses revealed no genetic partitioning for all species, indicating the Nile outflow does not currently, and may not have historically, posed a significant barrier to larval dispersal in these goatfish despite a present-day temperature and salinity gradient along the Mediterranean coastline of Egypt.

## Introduction

In Egyptian waters, there are many anthropogenic influences that can affect the genetic structure and diversity of marine aquatic organisms, such as those imposed by the Suez Canal migration corridor and the Nile Delta development projects. The Suez Canal opened in 1869, connects the Mediterranean Sea and the Red Sea, and provides a shorter shipping route between the Atlantic Ocean and the Indian Ocean. This conduit has allowed fish migration and colonization between both seas (e.g. Golani, 1998; Por, 2012; Steinitz, 1967). On the other hand, after the construction of the Aswan High Dam in Egypt between 1960 and 1970, the freshwater flow into the Mediterranean Sea from the mouth of the Nile River was significantly reduced. Moreover, 85 to 90% of the Egyptian population occupy the Nile Delta and areas adjacent to the Nile River, and this population has grown rapidly in the last two decades (Ali & El-Magd, 2016; Coleman *et al*., 2003; Embabi, 2018; Stanley & Warne, 1993). Thus, the Nile River Delta is no longer an active delta, but a wave-dominated coastal plain, with most of its former freshwater outflow diverted by a network of channels to irrigate Egyptian farmland.

In both the Mediterranean Sea and Red Sea, several studies have focused on the biology, fisheries management, genetics, and analytical chemistry of three species of goatfish (*M. barbatus, M. surmuletus, U. moluccensis*) because of their high economic value (Doğan & Ertan, 2017; Golani & Ritte, 1999; Keskin & Atar, 2013; Knebelsberger *et al*., 2014; Landi *et al*., 2014; Maggio *et al*., 2009; Mamuris *et al*., 1998; Mamuris *et al*., 2001; Matic-Skoko *et al*., 2018; Mehanna, 2009). However, no study to date has comprehensively examined patterns of connectivity using population genetic approaches of these three species.

Here we examine if there are concordant patterns of genetic structure for these three species of goatfish across this purported environmentally heterogeneous region. Comparative phylogeography has been used in many terrestrial and freshwater environments for delimiting regional phylogeographic patterns (Avise, 1996; Bowen *et al*., 2016; Hewitt, 1996), but few have considered marine habitat. We focused on three goatfish species from two different genera to test their genetic structure in the Nile Delta’s unique environment at the Mediterranean Sea’s periphery in Egypt.

## Materials and methods

Specimens of three species of goatfish (*Mullus barbatus* Linnaeus, 1758, *Mullus surmuletus* Linnaeus, 1758, and *Upeneus moluccensis* (Bleeker, 1855)) were obtained in August 2016 from the east (31°26’53.6”N 32°06’16.4”E) and west (31°04’24.7”N 29°02’21.7”E) coast of the Nile Delta in the Mediterranean Sea, near Alexandria, Egypt (Table 1 and Fig. 1). The specimens of goatfish were procured opportunistically from artisanal fishermen at landing sites and thus there were no requirements for the care and use of experimental animals or for collection permits from Egyptian authorities. Fish were identified by their morphological and meristic characters (Kim, 2002; Randall, 2001). All individuals were transported on ice slurries to the Fisheries Biology Laboratory of the National Institute of Oceanography and Fisheries, Alexandria Branch. Muscle biopsies and fin clips were preserved in ethanol (95%) for the population genetics study.

**Table 1.** Sample size and molecular diversity indices for three species of goatfish sampled to the west and east of Alexandria City, Egypt based on mitochondrial cytochrome c oxidase subunit I (COI) and cytochrome *b* (cyt b) sequence data.

| Gene, species, collection locality | Fragment size (bp) | $N^b$ | $H_N$ | Haplotype diversity<br>($h \pm SD$) | Nucleotide diversity<br>($\pi \pm SD$) | Fu's $F_S$ |
| --- | --- | --- | --- | --- | --- | --- |
| COI |  |  |  |  |  |  |
| *Mullus barbatus* |  |  |  |  |  |  |
| West Alexandria, Egypt | 655 | 19 | 8 | $0.82 \pm 0.06$ | $0.00304 \pm 0.00200$ | -2.37 |
| East Alexandria, Egypt | 655 | 24 | 6 | $0.67 \pm 0.07$ | $0.00201 \pm 0.00145$ | -1.17 |
| All samples | | 43 | 12 | $0.73 \pm 0.05$ | $0.00245 \pm 0.00166$ | **-5.47<sup>a</sup>** |
| *Mullus surmuletus* |  |  |  |  |  |  |
| West Alexandria, Egypt | 605 | 9 | 3 | $0.67 \pm 0.10$ | $0.00221 \pm 0.00171$ | 0.91 |
| East Alexandria, Egypt | 605 | 11 | 7 | $0.91 \pm 0.07$ | $0.00387 \pm 0.00258$ | -2.40 |
| All samples | | 20 | 7 | $0.79 \pm 0.06$ | $0.00302 \pm 0.00203$ | -1.49 |
| *Upeneus moluccensis* |  |  |  |  |  |  |
| West Alexandria, Egypt | 591 | 20 | 5 | $0.37 \pm 0.14$ | $0.00119 \pm 0.00105$ | -2.05 |
| East Alexandria, Egypt | 591 | 24 | 7 | $0.45 \pm 0.13$ | $0.00098 \pm 0.00091$ | **-5.40** |
| All samples | | 44 | 8 | $0.40 \pm 0.09$ | $0.00106 \pm 0.00094$ | **-5.39** |
| cyt *b* |  |  |  |  |  |  |
| *Mullus barbatus* |  |  |  |  |  |  |
| West Alexandria, Egypt | 401 | 24 | 3 | $0.52 \pm 0.07$ | $0.00268 \pm 0.00204$ | 1.55 |
| East Alexandria, Egypt | 401 | 24 | 6 | $0.70 \pm 0.06$ | $0.00317 \pm 0.00230$ | -1.29 |
| All samples | | 48 | 7 | $0.61 \pm 0.05$ | $0.00288 \pm 0.00210$ | -1.58 |
| *Mullus surmuletus* |  |  |  |  |  |  |
| West Alexandria, Egypt | 392 | 24 | 11 | $0.87 \pm 0.04$ | $0.00510 \pm 0.00332$ | **-5.21** |
| East Alexandria, Egypt | 392 | 23 | 4 | $0.66 \pm 0.06$ | $0.00301 \pm 0.00224$ | 0.52 |
| All samples | | 47 | 12 | $0.77 \pm 0.04$ | $0.00402 \pm 0.00270$ | **-5.39** |
| *Upeneus moluccensis* |  |  |  |  |  |  |
| West Alexandria, Egypt | 452 | 20 | 8 | $0.82 \pm 0.07$ | $0.00410 \pm 0.00274$ | -2.49 |
| East Alexandria, Egypt | 452 | 24 | 6 | $0.44 \pm 0.12$ | $0.00160 \pm 0.00138$ | **-3.01** |
| All samples | | 44 | 11 | $0.64 \pm 0.08$ | $0.00280 \pm 0.00200$ | **-5.72** |

<sup>a</sup>Numbers in bold are significant, $P < 0.02$ (Fu, 1997). <sup>b</sup>Abbreviations are as follows: $N$ , sample size; $H_N$ , number of haplotypes.

**Fig. 1.**
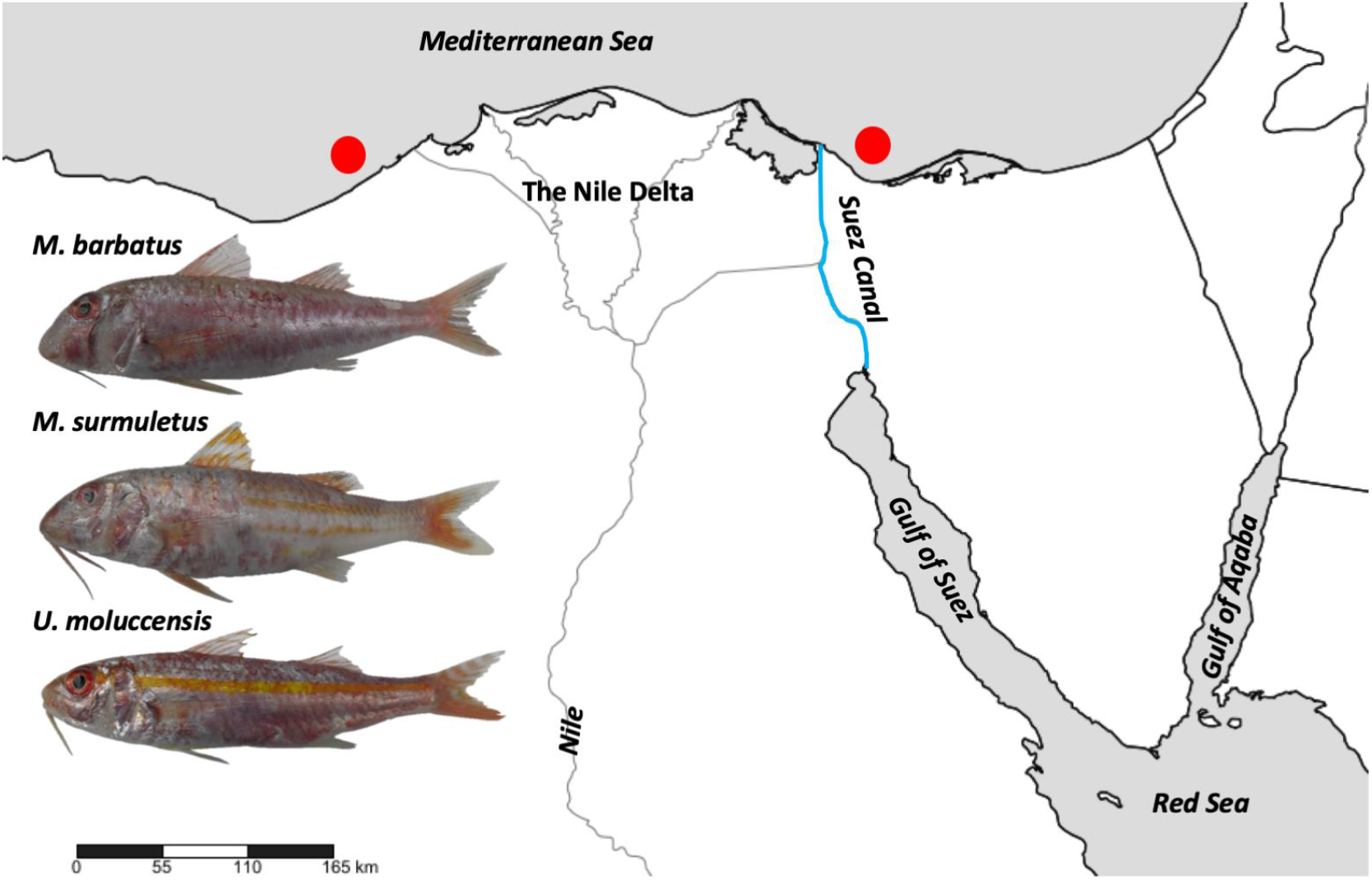
Map of sample collection sites to the west and east of Alexandria City, Egypt (red circles) for three species of goatfish (*Mullus barbatus, Mullus surmuletus*, and *Upeneus moluccensis*; photo credit: R. M. Fahim).

A DNeasy Blood & Tissue kit (Qiagen, Tokyo, Japan) was used to extract the genomic DNA from muscle tissue (∼25 mg) following the manufacturer’s protocol. Two primer sets of mtDNA [Cytochrome B gene (cyt b) L14724 5′ CGAAGCTTGATATGAAAAACCATCGTTG-3′ and H15149 5′-AAACTGCAGCCCCTCAGAATGATATTTGTCCTCA-3′ (Irwin *et al*., 1991) and Cytochrome Oxidase Subunit I (COI) FishF1, 5′-TCAACCAACCACAAAGACATTGGCAC-3′ and FishR1, 5′-TAGACTTCTGGGTGGCCAAAGAATCA-3′ (Ward *et al*., 2005)] were used for PCR amplification. PCR was performed with HotStarTaq Master Mix (Qiagen, Tokyo, Japan) in a 20 μl total reaction volume. The detailed PCR cycling conditions followed those outlined in previous reports (Soliman *et al*., 2017a, b). PCR amplification was tested using gel electrophoresis (1.5% agarose) and products that successfully amplified were cleaned with Exonuclease I and Shrimp Alkaline Phosphatase (Takara, Japan). The cleaned PCR products were cycle-sequenced using a BigDye™ Terminator v3.1 Cycle Sequencing Kit (ThermoFisher Scientific) following the manufacturer’s protocol. Both forward and reverse amplicons were sequenced using a Genetic Analyzer 3730xl (Applied Biosystems) at the Okinawa Institute of Science and Technology, Japan (OIST).

Sequences from both genes were edited and aligned using Geneious 8.1.9 (also see Table 1). Mitochondrial haplotype sequences generated in this study were deposited in GenBank (accession numbers: MG763621-MG763677). COI sequences from all three species were extracted from GenBank and added to the alignment (Appendix Table A1). Arlequin 3.5.1.2 (Excoffier *et al*., 2005) was used to calculate haplotype (*h*) and nucleotide diversity (*π*), as well as to test for population structure. jModelTest 1.0.1 (Posada, 2008) was used to select the best nucleotide substitution model using the Akaike information criterion (*AIC*). Genetic differentiation among sampling sites was first estimated with analysis of molecular variance (AMOVA) based on pairwise comparisons of sample groups; deviations from null distributions were tested with non-parametric permutation procedures (*N* = 99,999). Pairwise *Φ*_*ST*_ statistics were also calculated in Arlequin and significance tested by permutation (*N* = 99,999). Evolutionary relationships among haplotypes were evaluated by constructing median joining spanning networks (Bandelt *et al*., 1999) in PopART 1.7 (Leigh & Bryant, 2015). Deviations from neutrality were assessed with Fu’s *F*_*S*_ (Fu, 1997) for each species using Arlequin; significance was tested with 99,999 permutations.

## Results

Mitochondrial sequences from the three species of goatfish (*M. barbatus, M. surmuletus, U. moluccensis*) sampled to the west and east of Alexandria City on the Egyptian coastline, included 3 to 8 COI haplotypes and 3 to 11 cyt b haplotypes. Haplotype (*h*) and nucleotide (*π*) diversity ranged from 0.37 to 0.82 and 0.00098 to 0.00387 for COI, respectively, and 0.44 to 0.87 and 0.00160 to 0.00510 for cyt b (Table 1), respectively. Genetic diversity was not consistently higher in the west versus the east for any of the species (Table 1). Neutrality tests revealed negative and significant Fu’s *F*_*S*_ values in all samples of *M. barbatus* (−5.47) and *U. moluccensis* (−5.39) for COI, and *M. surmuletus* (−5.39) and *U. moluccensis* (−5.72) for cyt b (Table 1), but this pattern was not consistent between molecular markers.

The most common and second most common haplotypes in the median-joining spanning networks for each species were shared between individuals sampled to the west and east of Alexandria City for both COI (Fig. 2 A-C) and cyt b (Fig. 2 D-F). In addition, singleton haplotypes were not biased by geography (Fig. 2 A-F). AMOVA (Appendix Table A2) revealed no significant genetic structure for all three goatfish species for COI (*Φ*_*ST*_ = 0 to 0.013) and cyt b (*Φ*_*ST*_ = 0 to 0.045). The inclusion of supplementary COI sequences extracted from GenBank ranging as far north as the North Sea (for *M. surmuletus*), as far west as Portugal (for *M. barbatus* and *M. surmuletus*), and as far east as Taiwan (for *U. moluccensis*) did not provide evidence for cryptic lineages across broader geographic ranges based on the extensive sharing of common haplotypes (Fig. 2 G-I).

**Fig. 2.**
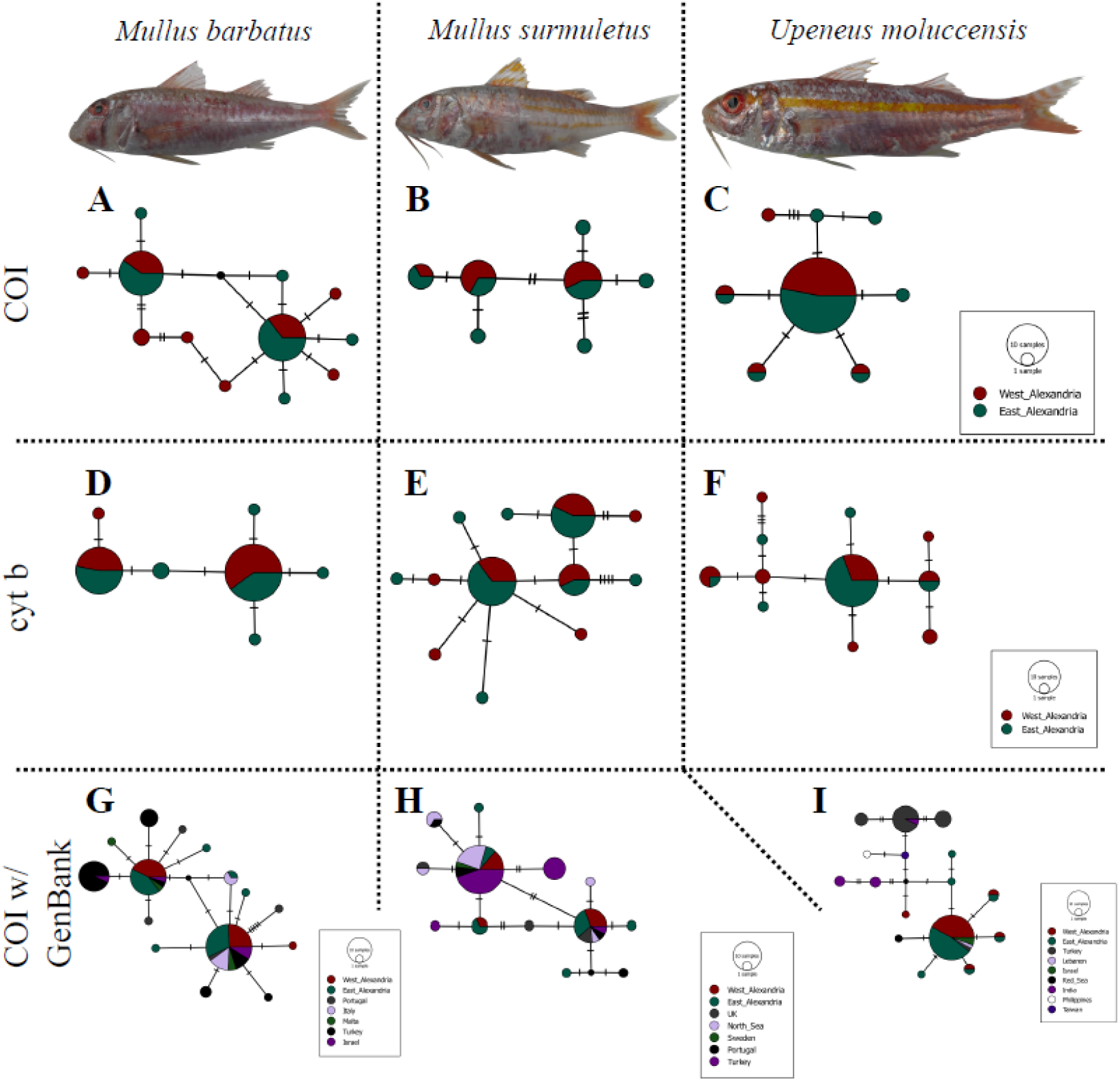
Median-joining networks showing relationships among mitochondrial cytochrome *c* oxidase subunit I (COI; A, B, C) and cytochrome *b* (cyt b; D, E, F) sequences for three species of goatfish (*Mullus barbatus*: A, D, G; *Mullus surmuletus*: B, E, H; *Upeneus moluccensis*: C, F, I) sampled to the west and east of Alexandria City, Egypt. COI and cyt b fragment length and site-specific samples sizes are provided in Table 1. Each circle represents a unique haplotype and its size is proportional to its total frequency (i.e. number of samples) as per the legend. Branches and black crossbars represent a single nucleotide change; colors denote collection location as indicated by the key. Supplementary COI data extracted from GenBank are considered in G, H, I (fragment length: 551 bp, 508 bp, 510 bp, respectively), and given in Supplementary Material Table S1 (photo credit: R. M. Fahim).

## Discussion

Our two marker (COI and cyt b) mtDNA dataset revealed that the genetic diversity was relatively high for *M. barbatus* and *M. surmuletus*. This result agrees with previous studies of population genetic structure using microsatellite markers for other fishes in the Mediterranean Sea (Félix-Hackradt *et al*., 2013; Galarza *et al*., 2009). Additionally, our results showed that the genetic variability in *M. surmuletus* was relatively higher than that in *M. barbatus* at both markers, which is supported by comparable RFLP results reported in Mamuris *et al*., (2001). In contrast, Matic-Skoko *et al*., (2018) used thirteen nuclear microsatellite markers for two of these goatfish species and found that the genetic diversity of *M. barbatus* was higher than *M. surmuletus* among 14 populations across the Mediterranean Sea.

The Lessepsian migrant species *U. moluccensis* showed the lowest genetic diversity among the three species of goatfishes considered in our study based on the COI marker but not the cyt b marker, possibly due to greater variability in the latter DNA fragment. A number of other fish species (e.g. *Dicentrarchus labrax, Diplodus sargus, Diplodus vulgaris*, and *Coptodon zillii*) demonstrated lower genetic diversity within the Mediterranean compared to populations outside of the Mediterranean Sea. Previous research also demonstrated significant genetic structure of *U. moluccensis* between the Red Sea and Mediterranean Sea (Golani & Ritte, 1999; Hassan & Bonhomme, 2005), which is congruent with the divergent morphology and stock assessments of this species from these two seas (Pazhayamadom *et al*., 2017). In our case, the COI haplotype network with supplemental GenBank sequences for this species did not provide evidence for significant differentiation across its Indo-West Pacific range, although we do note fixed differences for some locations (e.g. Red Sea, Taiwan, or Philippines), more so than any of the other goatfish species considered in our study. Significant genetic differentiation was additionally reported in range-wide studies of different goatfish species such as *Parupeneus multifasciatus* (Szabó *et al*., 2014) and *Mulloidichthys flavolineatus* (Fernandez-Silva *et al*., 2015), indicating cryptic lineages, something that we did not observe for any of the three species in our study. Congruence in genetic structure across different species, however, is seldom the case in comparative phylogeographic studies of reef organisms (Borsa *et al*., 2016; DiBattista *et al*., 2017; Lessios & Robertson, 2006; Selkoe *et al*., 2014; Tenggardjaja *et al*., 2016; Toonen *et al*., 2011).

Some studies have focused on genetic differentiation of marine and brackish populations using the COI gene within the northern Delta lakes and the Nile Delta in Egyptian waters (El-Nabi *et al*., 2017; Soliman *et al*., 2017; Abbas, 2018; Ali, 2019). It has been proposed that the construction of the Aswan Dam in the 1960s diminished the strength of the low salinity barrier due to freshwater outflow present prior to its construction (Vörösmarty & Sahagian, 2000; Alber, 2002). Higher relative salinity in this region following construction is also thought to have facilitated the recent colonization of the Mediterranean Sea by Red Sea invasive species via the Suez Canal (i.e. Lessepsian invaders; see Bernardi *et al*., 2016), which includes *U. moluccensis*. The ecological differences between the Red Sea and Mediterranean Sea might explain the genetic variance of invader goatfish species from the Red Sea to the Mediterranean Sea through the Suez Canal (Ahmed *et al*., 2016). However, some Lessepsian species might have been introduced from the Red Sea by alternative pathways, such as anthropogenic transport (e.g. Bariche *et al*., 2015). Thus, it is highly possible that signatures of genetic differentiation between the east and west Nile River Delta may no longer be apparent since the construction of the dam in the 1960s, particularly given that these goatfish species can tolerate varying levels of salinity. Indeed, the adult fish are often found in brackish waters (Hamza, 2009) and the differences in mean salinity across our study region was marginal (∼3.5 parts per thousand difference; see Fig. 3b). Such information suggests that goatfish species may have been tolerant to any original freshwater barrier that existed before dam construction. Thus, given the tolerance of *Mullus* goatfish to brackish waters, the most parsimonious explanation is that no such genetic break existed in the past. Current day gradients based on sea surface temperature also suggest the presence of only a marginal gradient across our study region (∼0.3 °C difference; see Fig. 3a), and so it too likely poses minimal disruption to ongoing gene flow. As an alternative explanation, it is possible the molecular markers utilized in this study may not have the resolution to detect genetic breaks (e.g. Bi *et al*., 2009), and so future research would be well-served by incorporating finer-scale molecular methodologies such as microsatellites (e.g. Félix-Hackradt *et al*., 2013; Galarza *et al*., 2009) or single-nucleotide polymorphisms (SNPs; Martinsohn *et al*., 2009).

**Fig. 3.**
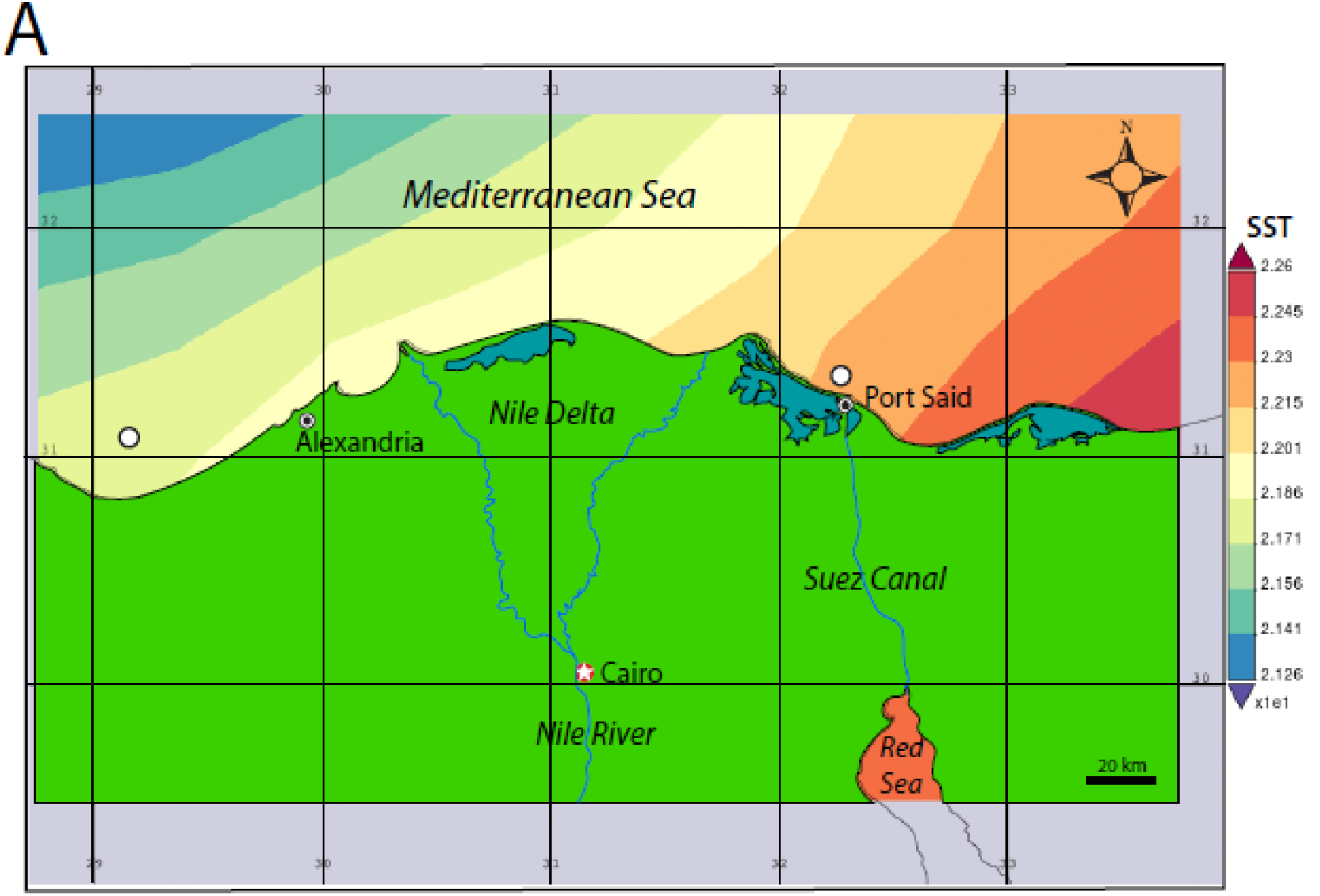
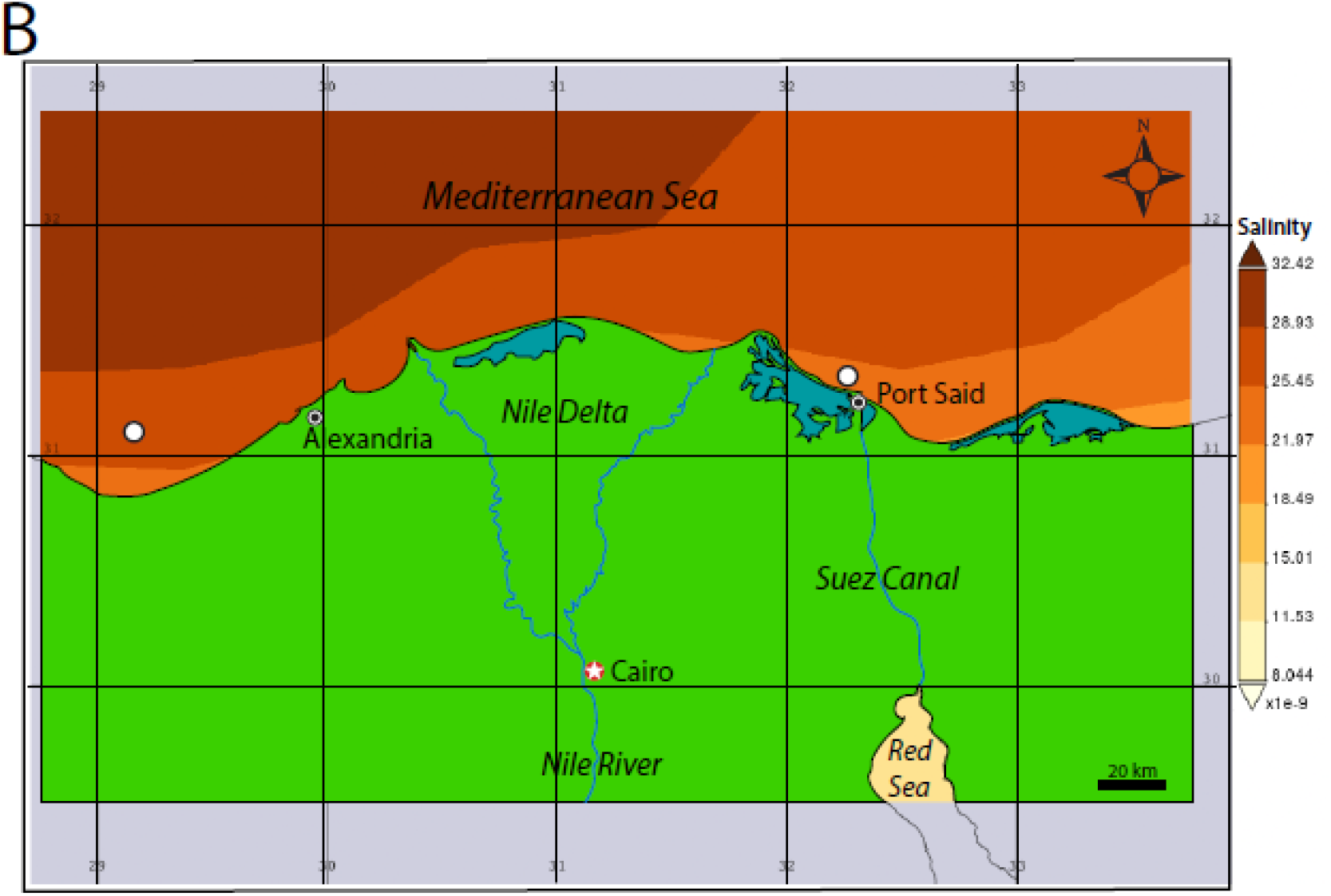
Maps of sample collection sites (empty white circles) and ocean regions overlaid with A) sea surface temperature (SST in °C) and B) salinity (in parts per thousand) estimates produced with the Giovanni online data system, which was developed and maintained by the NASA GES DISC.

## Acknowledgments

We are grateful to Prof. Gordon Arbuthnott (OIST) for his support in providing chemicals and sequencing facilities. Analyses and visualizations used in this paper based on environmental parameters were produced with the Giovanni online data system, developed and maintained by the NASA GES DISC. This research was supported by an internal grant of the Okinawa Institute of Science and Technology (OIST), Japan to the research group from the Faculty Affairs Office, and a Curtin University Early Career Research Fellowship (ECRF) to J.D.D.

