## Appendix Table A1 for "Preliminary genetic assessment of three goatfish species in the Mediterranean Sea"

**Appendix Table A1** Origin of supplemental mitochondrial cytochrome c oxidase subunit I (COI) sequences included in this study from NCBI.

| tissue ID | species | sample location | publication | photo available | GenBank accession # |
| --- | --- | --- | --- | --- | --- |
| TR054EK | *Mullus barbatus* | Turkey | unpublished | no | JQ623957 |
| FCFOPS155 | *Mullus barbatus* | Portugal, Faro, Algarve, Sub-area IXa | Costa *et al.,* 2012**^a^** | no | JQ774687 |
| TR999EK | *Mullus barbatus* | Turkey | Keskin & Atar, 2013 | no | KC500953 |
| TR997EK | *Mullus barbatus* | Turkey | Keskin & Atar, 2013 | no | KC500955 |
| TR996EK | *Mullus barbatus* | Turkey | Keskin & Atar, 2013 | no | KC500956 |
| TR995EK | *Mullus barbatus* | Turkey | Keskin & Atar, 2013 | no | KC500957 |
| TR994EK | *Mullus barbatus* | Turkey | Keskin & Atar, 2013 | no | KC500958 |
| TR993EK | *Mullus barbatus* | Turkey | Keskin & Atar, 2013 | no | KC500959 |
| TR992EK | *Mullus barbatus* | Turkey | Keskin & Atar, 2013 | no | KC500960 |
| TR991EK | *Mullus barbatus* | Turkey | Keskin & Atar, 2013 | no | KC500961 |
| TR990EK | *Mullus barbatus* | Turkey | Keskin & Atar, 2013 | no | KC500962 |
| TR989EK | *Mullus barbatus* | Turkey | Keskin & Atar, 2013 | no | KC500963 |
| TR988EK | *Mullus barbatus* | Turkey | Keskin & Atar, 2013 | no | KC500964 |
| TR987EK | *Mullus barbatus* | Turkey | Keskin & Atar, 2013 | no | KC500965 |
| TR986EK | *Mullus barbatus* | Turkey | Keskin & Atar, 2013 | no | KC500966 |
| TR984EK | *Mullus barbatus* | Turkey | Keskin & Atar, 2013 | no | KC500968 |
| TR983EK | *Mullus barbatus* | Turkey | Keskin & Atar, 2013 | no | KC500969 |
| TR982EK | *Mullus barbatus* | Turkey | Keskin & Atar, 2013 | no | KC500970 |
| TR981EK | *Mullus barbatus* | Turkey | Keskin & Atar, 2013 | no | KC500971 |
| TR980EK | *Mullus barbatus* | Turkey | Keskin & Atar, 2013 | no | KC500972 |
| CSFOM-126 | *Mullus barbatus* | Italy, Sicily | Landi *et al.,* 2014 | no | KJ709562 |
| CSFOM-063 | *Mullus barbatus* | Italy, Sicily | Landi *et al.,* 2014 | no | KJ709563 |
| CSFOM-062 | *Mullus barbatus* | Italy, Sicily | Landi *et al.,* 2014 | no | KJ709564 |
| CSFOM-125 | *Mullus barbatus* | Italy, Sicily | Landi *et al.,* 2014 | no | KJ709565 |
| CSFOM-234 | *Mullus barbatus* | Italy, Sicily | Landi *et al.,* 2014 | no | KJ709566 |
| CSFOM-127 | *Mullus barbatus* | Italy, Sicily | Landi *et al.,* 2014 | no | KJ709567 |
| CSFOM-128 | *Mullus barbatus* | Italy, Sicily | Landi *et al.,* 2014 | no | KJ709568 |
| MCFS06028 | *Mullus barbatus* | Malta (36.233° N 14.9° E) | Landi *et al.,* 2014 | no | KJ709823 |
| MCFS06022 | *Mullus barbatus* | Malta (36.233° N 14.9° E) | Landi *et al.,* 2014 | no | KJ709824 |
| MCFS06071 | *Mullus barbatus* | Malta (35.567° N 15.267° E) | Landi *et al.,* 2014 | no | KJ709825 |
| MCFS06024 | *Mullus barbatus* | Malta (36.233° N 14.9° E) | Landi *et al.,* 2014 | no | KJ709827 |
| MLFPI255 | *Mullus barbatus* | Portugal, Algarve (37.075° N 8.5033° W) | Landi *et al.,* 2014 | yes | KJ768261 |
| MLFPI201 | *Mullus barbatus* | Portugal, Algarve (37.947° N 9.995° W) | Landi *et al.,* 2014 | yes | KJ768262 |
| MLFPI254 | *Mullus barbatus* | Portugal, Algarve (37.075° N 8.5033° W) | Landi *et al.,* 2014 | yes | KJ768263 |
| MuBa29G | *Mullus barbatus* | Israel | unpublished | yes | KM538414 |
| MuBa29L | *Mullus barbatus* | Israel | unpublished | yes | KM538415 |
| MuBa29K | *Mullus barbatus* | Israel | unpublished | yes | KM538416 |
| MuBa29J | *Mullus barbatus* | Israel | unpublished | yes | KM538417 |
| MuBa29I | *Mullus barbatus* | Israel | unpublished | yes | KM538418 |
| Bsex-320 | *Mullus barbatus* | Turkey, Black Sea | unpublished | no | KP136668 |
| Bsex-321 | *Mullus barbatus* | Turkey, Black Sea | unpublished | no | KP136669 |
| Bsex-75 | *Mullus barbatus* | Turkey, Black Sea | unpublished | no | KP136730 |
| Bsex-76 | *Mullus barbatus* | Turkey, Black Sea | unpublished | no | KP136731 |
| Bsex-77 | *Mullus barbatus* | Turkey, Black Sea | unpublished | no | KP136732 |
| Bsex-78 | *Mullus barbatus* | Turkey, Black Sea | unpublished | no | KP136733 |
| Bsex-79 | *Mullus barbatus* | Turkey, Black Sea | unpublished | no | KP136734 |
| 454 | *Mullus barbatus* | Turkey, Istanbul, Silivri | unpublished | no | KY176532 |
| FCFOP64-39 | *Mullus surmuletus* | Portugal (38.22° N 8.83° W) | Costa *et al.,* 2012 | yes | JQ774872 |
| FCFOP64-41 | *Mullus surmuletus* | Portugal (38.22° N 8.83° W) | Costa *et al.,* 2012 | yes | JQ774873 |
| FCFOP64-43 | *Mullus surmuletus* | Portugal (38.22° N 8.83° W) | Costa *et al.,* 2012 | yes | JQ774874 |
| FCFOP64-42 | *Mullus surmuletus* | Portugal (38.22° N 8.83° W) | Costa *et al.,* 2012 | yes | JQ774875 |
| FCFOP64-40 | *Mullus surmuletus* | Portugal (38.22° N 8.83° W) | Costa *et al.,* 2012 | yes | JQ774876 |
| TR1004EK | *Mullus surmuletus* | Turkey | Keskin & Atar, 2013 | no | KC500973 |
| TR1006EK | *Mullus surmuletus* | Turkey | Keskin & Atar, 2013 | no | KC500974 |
| TR1007EK | *Mullus surmuletus* | Turkey | Keskin & Atar, 2013 | no | KC500975 |
| TR1008EK | *Mullus surmuletus* | Turkey | Keskin & Atar, 2013 | no | KC500976 |
| TR1009EK | *Mullus surmuletus* | Turkey | Keskin & Atar, 2013 | no | KC500977 |
| TR1010EK | *Mullus surmuletus* | Turkey | Keskin & Atar, 2013 | no | KC500978 |
| TR1011EK | *Mullus surmuletus* | Turkey | Keskin & Atar, 2013 | no | KC500979 |
| TR1012EK | *Mullus surmuletus* | Turkey | Keskin & Atar, 2013 | no | KC500980 |
| TR1013EK | *Mullus surmuletus* | Turkey | Keskin & Atar, 2013 | no | KC500981 |
| TR1014EK | *Mullus surmuletus* | Turkey | Keskin & Atar, 2013 | no | KC500982 |
| TR1015EK | *Mullus surmuletus* | Turkey | Keskin & Atar, 2013 | no | KC500983 |
| TR1016EK | *Mullus surmuletus* | Turkey | Keskin & Atar, 2013 | no | KC500984 |
| TR1017EK | *Mullus surmuletus* | Turkey | Keskin & Atar, 2013 | no | KC500985 |
| TR1019EK | *Mullus surmuletus* | Turkey | Keskin & Atar, 2013 | no | KC500987 |
| TR1003EK | *Mullus surmuletus* | Turkey | Keskin & Atar, 2013 | no | KC500988 |
| TR1005EK | *Mullus surmuletus* | Turkey | Keskin & Atar, 2013 | no | KC500989 |
| TR1000EK | *Mullus surmuletus* | Turkey | Keskin & Atar, 2013 | no | KC500990 |
| TR1001EK | *Mullus surmuletus* | Turkey | Keskin & Atar, 2013 | no | KC500991 |
| TR1002EK | *Mullus surmuletus* | Turkey | Keskin & Atar, 2013 | no | KC500992 |
| NRM:50181 | *Mullus surmuletus* | Sweden, Skagerrak | unpublished | no | KJ128554 |
| MT04167 | *Mullus surmuletus* | North Sea (58.325° N 1.328° W) | Knebelsberger *et al.,* 2014 | yes | KJ205067 |
| MT01843 | *Mullus surmuletus* | North Sea, German Bight (54.5° N 7° E) | Knebelsberger *et al.,* 2014 | yes | KJ205068 |
| MT04165 | *Mullus surmuletus* | North Sea (59.147° N 1.109° W) | Knebelsberger *et al.,* 2014 | yes | KJ205070 |
| MT01844 | *Mullus surmuletus* | North Sea, German Bight (54.5° N 7° E) | Knebelsberger *et al.,* 2014 | yes | KJ205071 |
| MT02895 | *Mullus surmuletus* | North Sea (53.862° N 4.617° E) | Knebelsberger *et al.,* 2014 | yes | KJ205072 |
| MT02284 | *Mullus surmuletus* | North Sea (58.802° N 0.602° W) | Knebelsberger *et al.,* 2014 | yes | KJ205073 |
| MT01846 | *Mullus surmuletus* | North Sea, German Bight (54.5° N 7° E) | Knebelsberger *et al.,* 2014 | yes | KJ205074 |
| MT01847 | *Mullus surmuletus* | North Sea, German Bight (54.5° N 7° E) | Knebelsberger *et al.,* 2014 | yes | KJ205076 |
| MT02295 | *Mullus surmuletus* | North Sea (58.871° N 2.322° W) | Knebelsberger *et al.,* 2014 | yes | KJ205077 |
| MT02938 | *Mullus surmuletus* | North Sea (55.343° N 3.699° E) | Knebelsberger *et al.,* 2014 | yes | KJ205079 |
| MT02985 | *Mullus surmuletus* | North Sea (54.37° N 7.088° E) | Knebelsberger *et al.,* 2014 | yes | KJ205080 |
| MT04164 | *Mullus surmuletus* | North Sea (59.763° N 0.372° E) | Knebelsberger *et al.,* 2014 | yes | KJ205081 |
| DWCS06-136 | *Mullus surmuletus* | United Kingdom, Bristol Channel, VIIf (51.36° N 4.23° W) | Knebelsberger *et al.,* 2014 | no | KJ205292 |
| DWCS06-101 | *Mullus surmuletus* | United Kingdom, Bristol Channel, VIIf (51.36° N 4.34° W) | Knebelsberger *et al.,* 2014 | no | KJ205293 |
| DWCS06-156 | *Mullus surmuletus* | United Kingdom, Bristol Channel, VIIf (51.32° N 4.31° W) | Knebelsberger *et al.,* 2014 | no | KJ205294 |
| DWCS06-103 | *Mullus surmuletus* | United Kingdom, Bristol Channel, VIIf (51.36° N 4.34° W) | Knebelsberger *et al.,* 2014 | no | KJ205295 |
| 1749 | *Mullus surmuletus* | Turkey: Gulf of Iskenderun | unpublished | no | KY176534 |
| 1752 | *Mullus surmuletus* | Turkey: Gulf of Iskenderun | unpublished | no | KY176535 |
| INDAPKKD-46 | *Upeneus moluccensis* | India, Kakinada | unpublished | no | FJ265831 |
| TR111EK | *Upeneus moluccensis* | Turkey | unpublished | no | JQ624014 |
| TR1865EK | *Upeneus moluccensis* | Turkey | Keskin & Atar, 2013 | no | KC501833 |
| TR1860EK | *Upeneus moluccensis* | Turkey | Keskin & Atar, 2013 | no | KC501834 |
| TR1861EK | *Upeneus moluccensis* | Turkey | Keskin & Atar, 2013 | no | KC501835 |
| TR1862EK | *Upeneus moluccensis* | Turkey | Keskin & Atar, 2013 | no | KC501836 |
| TR1863EK | *Upeneus moluccensis* | Turkey | Keskin & Atar, 2013 | no | KC501837 |
| TR1864EK | *Upeneus moluccensis* | Turkey | Keskin & Atar, 2013 | no | KC501838 |
| TR1866EK | *Upeneus moluccensis* | Turkey | Keskin & Atar, 2013 | no | KC501839 |
| TR1879EK | *Upeneus moluccensis* | Turkey | Keskin & Atar, 2013 | no | KC501840 |
| TR1878EK | *Upeneus moluccensis* | Turkey | Keskin & Atar, 2013 | no | KC501841 |
| TR1877EK | *Upeneus moluccensis* | Turkey | Keskin & Atar, 2013 | no | KC501842 |
| TR1876EK | *Upeneus moluccensis* | Turkey | Keskin & Atar, 2013 | no | KC501843 |
| TR1867EK | *Upeneus moluccensis* | Turkey | Keskin & Atar, 2013 | no | KC501844 |
| TR1875EK | *Upeneus moluccensis* | Turkey | Keskin & Atar, 2013 | no | KC501845 |
| TR1874EK | *Upeneus moluccensis* | Turkey | Keskin & Atar, 2013 | no | KC501846 |
| TR1873EK | *Upeneus moluccensis* | Turkey | Keskin & Atar, 2013 | no | KC501847 |
| TR1872EK | *Upeneus moluccensis* | Turkey | Keskin & Atar, 2013 | no | KC501848 |
| TR1871EK | *Upeneus moluccensis* | Turkey | Keskin & Atar, 2013 | no | KC501849 |
| TR1869EK | *Upeneus moluccensis* | Turkey | Keskin & Atar, 2013 | no | KC501851 |
| TR1868EK | *Upeneus moluccensis* | Turkey | Keskin & Atar, 2013 | no | KC501852 |
| ARO 174 | *Upeneus moluccensis* | Philippines, Aurora, Region 3 | unpublished | no | KF009674 |
| CIFE:FGB UM-01 | *Upeneus moluccensis* | India (8.81° N 78.14° E) | unpublished | no | KJ920110 |
| CIFE:FGB UM-02 | *Upeneus moluccensis* | India (8.81° N 78.14° E) | unpublished | no | KJ920112 |
| CIFE:FGB UM-03 | *Upeneus moluccensis* | India (8.81° N 78.14° E) | unpublished | no | KM079325 |
| CIFE:FGB UM-04 | *Upeneus moluccensis* | India (8.81° N 78.14° E) | unpublished | no | KM079326 |
| UpMo40T | *Upeneus moluccensis* | Israel | unpublished | yes | KM538622 |
| UpMo40S | *Upeneus moluccensis* | Israel | unpublished | yes | KM538623 |
| AUBM Lot 50 | *Upeneus moluccensis* | Lebanon | unpublished | no | KR861567 |
| n/a | *Upeneus moluccensis* | Egypt, Red Sea | unpublished | no | KT864716 |
| ASIZP0801469 | *Upeneus moluccensis* | Taiwan | Chang *et al.,* 2016 | no | KU944148 |
| 1066 | *Upeneus moluccensis* | Turkey, Antalya, Side | unpublished | no | KY176689 |

**^a^**References: Chang *et al.,* 2016, *Molecular Ecology Resources,* 17, 796-805; Costa *et al.,* 2012, *PLoS ONE,* 7, E35858; Keskin & Atar, 2013, *Molecular Ecology Resources,* 13, 788-797; Knebelsberger *et al.,* 2014, *Molecular Ecology Resources,* 14, 1060-1071; Landi *et al.,* 2014, *PLoS ONE,* 9, E106135.
