## Appendix Table A2 for "Preliminary genetic assessment of three goatfish species in the Mediterranean Sea"

**Appendix Table A2** Matrix of population pairwise *Φ_ST_* values, with associated *P*-values in parentheses, based on mitochondrial DNA cytochrome c oxidase subunit I (COI) and cytochrome b (cyt b) sequences for three species of goatfish sampled to the west and east of Alexandria City, Egypt. COI and cyt b fragment length and site-specific samples sizes are provided in Table 1. All negative *Φ_ST_* values were adjusted to zero.

*Mullus barbatus*

| COI | | | cyt b | | |
| --- | --- | --- | --- | --- | --- |
| site | West | East | site | West | East |
| West | **-** |  | West | **-** |  |
| East | 0.013 (0.252) | **-** | East | 0 (0.879) | **-** |

*Mullus surmuletus*

| COI | | | cyt b | | |
| --- | --- | --- | --- | --- | --- |
| site | West | East | site | West | East |
| West | **-** |  | West | **-** |  |
| East | 0 (0.993) | **-** | East | 0 (0.961) | **-** |

*Upeneus moluccensis*

| COI | | | cyt b | | |
| --- | --- | --- | --- | --- | --- |
| site | West | East | site | West | East |
| West | **-** |  | West | **-** |  |
| East | 0 (0.962) | **-** | East | 0.045 (0.076) | **-** |
